# A bibliometric analysis of neurobiological and behavioral disturbances of cafeteria diet interventions

**DOI:** 10.1101/2024.01.09.574927

**Authors:** Alejandra Lopez-Castro

**Affiliations:** Instituto de Neurobiologia, Universidad Nacional Autonoma de Mexico campus Juriquilla, Queretaro, Mexico City

## Abstract

Obesity is a global epidemic mainly caused by the overconsumption of western diets, high in fat and sugars. Cafeteria diet administered to rodents is an effective model of the metabolic, neurobiological, and behavioral disturbances caused by the over consumption of western diet in humans. However, this is still an emerging research field. To provide information about the past, present and future of the research field, this study aims to explore the research field of cafeteria diet and behavior through bibliometric analysis. Original articles on cafeteria diet and behavior were obtained from Pubmed, Scopus and Web of Science databases from 2013 to Octuber 30, 2023. The R packages litsearchr, bibliometrix, sjrdata and mblm were used for descriptive and inferential statistics. Linear regression, concept mapping and trend analysis were used for relationship analysis. 85 articles included from 457 authors, 20 countries and 56 institutions were included. 46 from Pubmed, 12 from Scopus and 27 from Web of Science. The 25 topmost productive authors were from Spain, Brazil, Australia, Switzerland, and USA. 15 authors had an h-index higher than 3. The institution with the largest production of articles is the University of South Wales with 10 articles. A simple linear regression could not establish significance between the relationship between the impact factor and the number of citations received. In addition, a conceptual structure map was performed, and 5 clusters were found. Finally, by a bi-factor analysis, a trend topic established that anxiety is the term currently in trend and since 2017 in the cafeteria diet and behavior research field. The present study explores the performance of authors, countries, institutions, and journals on classical measures of scientific parameters. This helped to model multiple correspondence and trend analyses that provide a reliable source of information to direct research on cafeteria diet interventions.

## Introduction

Obesity (BMI 30 kg/m^2^) has reached epidemic proportions and, according to the World Health Organization (WHO), by 2023 over 1 billion adults will have obesity (1). At the individual level, the WHO recommends reducing and limit energy intake from total fats and sugars (2). And at the population level, “*Evidence-based, impactful and cost-effective actions*” includes increasing the cost of sugar-sweetened beverages and reducing the sugar content of beverages. Still, there is no intervention on foods rich in both sugar and fats (known as Western diets), perhaps due to the lack of evidence from translational research.

One way to provide further evidence of the effect of Western diet consumption is the administration of Cafeteria Diet (CAF) to rodents. The CAF diet is composed of cookies, sausages, sugary juices, fried foods, and sweets commonly consumed by humans in their daily life context (3). Given its composition and efficacy to induce metabolic disturbances, the CAF diet is proposed as the ideal diet to study the consequences of junk food consumption (3). Altogether, this model allows us to understand the etiology of obesity and to develop effective approaches in the clinic (4).

The physiological effects of the CAF diet on body weight and metabolism have been widely described, resulting in increased adiposity, dyslipidemia, insulin resistance, and other complications related to metabolic syndrome (5). In addition, excessive consumption of CAF diet is the cause of hepatic, pancreatic, adipose, renal, and reproductive dysfunction(6).

CAF diet consumption has also been associated with negative neurobiological outcomes (7); (3). Also, CAF diet alters the microenvironment of several metabolic and non-metabolic brain regions (hypothalamus, hippocampus, amygdala, cerebral cortex, choroid plexus, etc.) impairing neuronal surveillance, synapses, plasticity and neuronal-glial communication (8). At the cellular level, CAF diet alters cell signaling and mitochondrial function, and increases oxidative stress an (10,11). This implies that neurobiological disturbances can be approached as early markers of obesity and its comorbidities (12) and need to be studied thoroughly (13).

Negative behavioral outcomes caused by CAF diet consumption have also been reported. For example, consumption of the CAF diet impairs cognition, memory, reward, and locomotion (14). Moreover, CAF diet alters emotional regulation, increasing stress-related disorders (15), anxiety-like behavior, depression-like behavior (16), and disrupted eating patterns (17). The underlying mechanisms and the relationship between the CAF diet and the behavioral factor is an emerging and growing field of research.

In health care, clinicians consider behavioral factors in overweight and obese patients to be more important than the biological factors (18), as claimed by the beneficial effect of cognitive behavior intervention on treatment adherence (19). In addition, cognitive-behavior wellness promotes better life quality, and better lifestyle changes in the long term (20). It is undeniable that obesity is a multifactorial problem and not just a metabolic problem. Therefore, it cannot be addressed by limited interventions. Thus, analyses of the CAF diet effects in animal models are necessary to direct research toward yet unanswered questions and new perspectives to address them (21).

Bibliographic analysis uses statistical tools, science mapping, and performance methodology to analyze and provide quantitative data on scientific publications. It may assess research performance, author productivity, and impact of manuscript (22). Thus, to help assess the course of cafeteria diet research and its relationship with behavior, in this study, we present an analysis of authors, citations, countries and trending topics from 2013 to 2023.

## Materials and methods

### 0.1 Search strategy and data source

This study is a bibliometric analysis with data obtained from Scopus, Pubmed and Web of Science (WOS) databases. Given that western diets are characterized by a high proportion of sugar and fats, reports with similar macronutrient composition as cafeteria diets (high-sugar and high-fat) were also considered. R Studio 2023.09.0 (23) and litsearchr 1.0.0 (24) package was used to automate search term selection and search strategy. A naïve search on the three databases (Table 1) was performed, with two central topics of interest (“cafeteria diet”) AND (behavior), limiting results to original articles from 2013 to 2023, and published in english. All three databases were merged into a single database where duplicated titles and abstracts were removed.

**Table 1.**
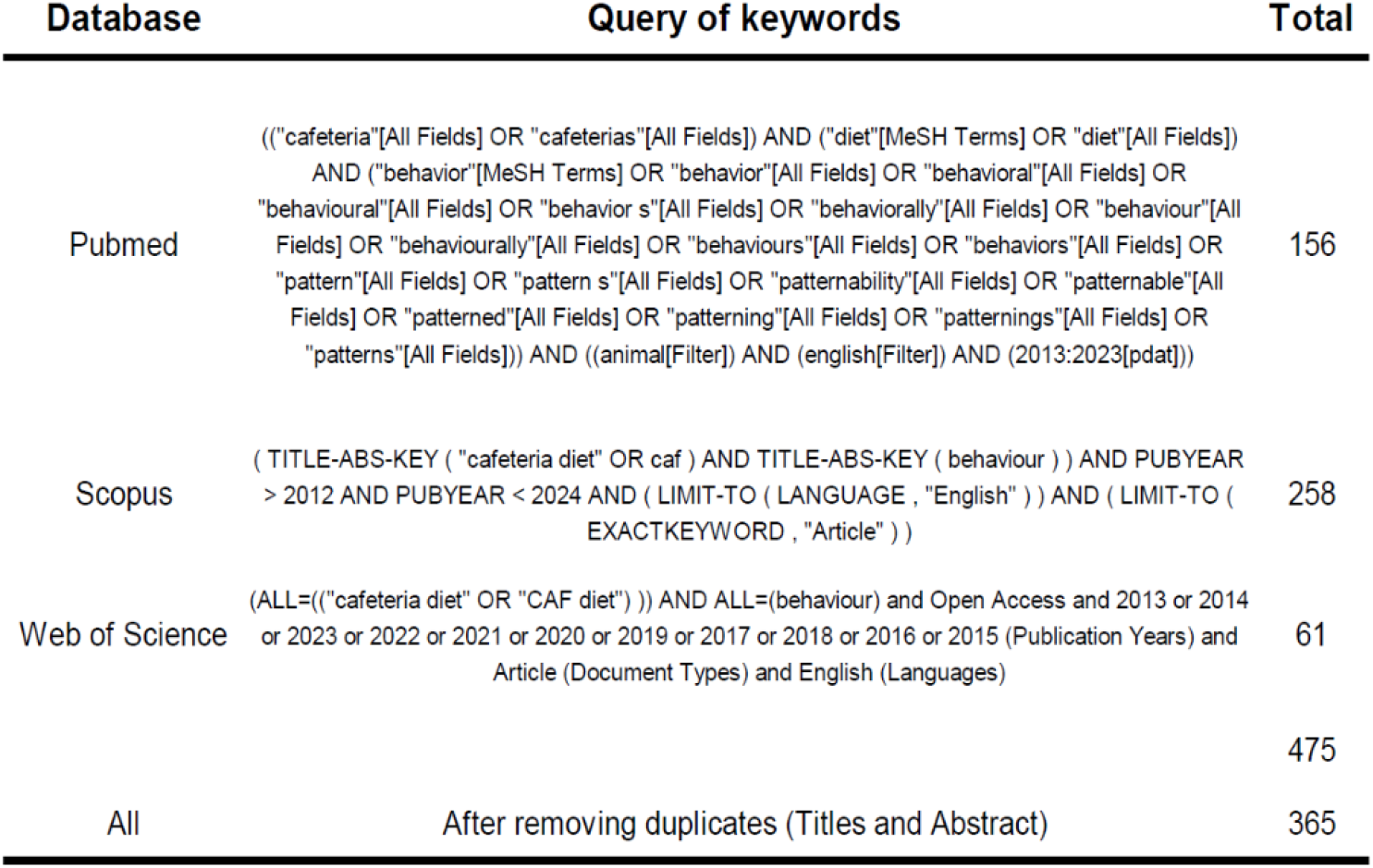
The Pubmeb search yielded 156 documents, 258 in Scopus and 61 in Web of Science (WOS) with a total of 475 articles that were filtered by removing duplicates in title and abstract, resulting in a total final of 365 documents.

| Database | Query of keywords | Total |
| --- | --- | --- |
| Pubmed | (("cafeteria"[All Fields] OR "cafeterias"[All Fields]) AND ("diet"[MeSH Terms] OR "diet"[All Fields]) AND ("behavior"[MeSH Terms] OR "behavior"[All Fields] OR "behavioral"[All Fields] OR "behavioural"[All Fields] OR "behavior s"[All Fields] OR "behaviorally"[All Fields] OR "behaviour"[All Fields] OR "behaviourally"[All Fields] OR "behaviours"[All Fields] OR "behaviors"[All Fields] OR "pattern"[All Fields] OR "pattern s"[All Fields] OR "patternability"[All Fields] OR "patternable"[All Fields] OR "patterned"[All Fields] OR "patterning"[All Fields] OR "patternings"[All Fields] OR "patterns"[All Fields])) AND ((animal[Filter]) AND (english[Filter]) AND (2013:2023[pdat])) | 156 |
| Scopus | ( TITLE-ABS-KEY ( "cafeteria diet" OR caf ) AND TITLE-ABS-KEY ( behaviour ) ) AND PUBYEAR > 2012 AND PUBYEAR < 2024 AND ( LIMIT-TO ( LANGUAGE , "English" ) ) AND ( LIMIT-TO ( EXACTKEYWORD , "Article" ) ) | 258 |
| Web of Science | (ALL=(("cafeteria diet" OR "CAF diet" ) ) AND ALL=(behaviour) and Open Access and 2013 or 2014 or 2023 or 2022 or 2021 or 2020 or 2019 or 2017 or 2018 or 2016 or 2015 (Publication Years) and Article (Document Types) and English (Languages) | 61 |
|  |  | 475 |
| All | After removing duplicates (Titles and Abstract) | 365 |

### 0.2 Eligibility criteria

Eligible studies were reviewed and only considered when *cafeteria diet* was associated with a) behavioral disturbances including cognitive impairments, stress, anxiety, depression-like behavior, and feeding behavior; and b) neurobiological disturbances including neuroinflammation, cell cycle and neurogenesis, metabolic and hormonal signaling, cell stress, and neural structure and function.

Studies were excluded when: 1) Had high-fat diets or western diets rich only in fats nor sugars; 2) were conducted in dams, during gestation or lactation; 3) used genetically modified rodents (e.g., transgenic, or obesity-prone). For this analysis were not considered conference articles, books, notes, editorials, brief surveys, letters, and book chapters. The data were completed with the help of SJR database from 2009 to 2022. A total of 116 articles were assessed for eligibility after eligibility criteria. Manual filtering of irrelevant articles was performed based on the title, abstract, and main text. A total of 85 articles (Table S1) were then included for the bibliometric analysis. This study followed the PRISMA-ScR checklist (Fig 1).

**Fig 1.**
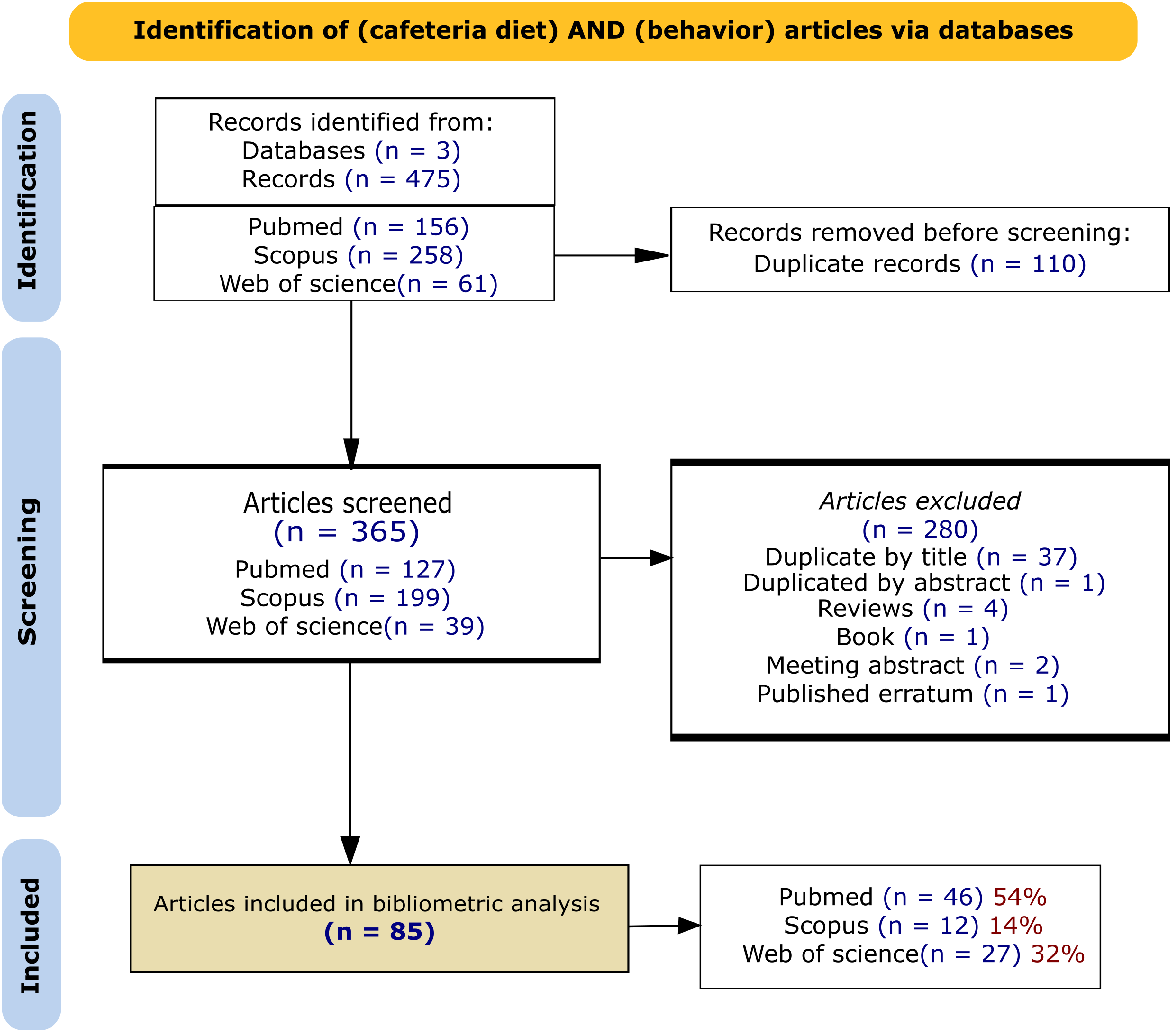
PRISMA-ScR. Flowchart of the search methodology and data source of this bibliometric analysis.

### 0.3 Data processing

The information extracted from the retrieved documents included: publication year, title, keywords, abstract, author, country, affiliation, journal and number of citations, impact factor (IF), publisher, categories, areas, SJR best quartile and H index. The Impact Factor (IF) for the selected journals was retrieved from JCRImpactFactor by Clarivate 1.0.0 package (25).

### 0.4 Data analysis

Bibliometric analysis was performed using R packages bibliometrix 4.1.3 (26), litsearchr (24), and sjrdata 0.4.0 were used. These libraries contain a series of functions for scientometric quantitative research and SCImago Journal & Country Rank Data (27). Two analysis methods were used: performance analysis (contributions of research constituents) and (2) science mapping (the relationships between research constituents); a detailed description of both techniques has been recently presented (28).

Performance analysis metrics reported are:

1. Publication-related metrics: total number of articles, annual growth rate (%), average citation per article, number of total authors, co-authored publications, and single-authored publications, international co-authorship, author’s keywords, and keywords provided by the publisher.
2. Citation-related metrics: total citations per author, number of publications per author and publication year. And the 10 most cited articles.
3. Citation and publication-related metrics: h-index, g-index, and m-index. The h-index is a measure indicating that the author has at least h citations that have h + 1 citations (29). The g-index is calculated as a combination of the articles that together have g^2^ citations (30). The m-index is calculated with the h-index divided by the number of years since their first published paper.

For the science mapping analyses, the following parameters were obtained:

1. Citation analysis: article by country and citations per country. Followed by production per institution and the analysis between impact factor and number of citations. For the latter, we explored the correlation of both variables with a Pearson correlation and then used a simple linear regression model.
2. Co-word analysis: Key terms were collected from authors and journals keywords, titles, and abstracts. To avoid stop words or uninformative words (general science or data analysis words), adverbs, articles and method descriptors were filtered. A document term-matrix (dfm) was performed to obtain the frequency and weight of terms appearing in the collection of articles (Table S2). To analyze the relationships between terms, the multiple correspondence analysis (MCA) method was applied. To facilitate later interpretation of the resulting groups and better identification of association patterns (Fig S1), k-means with 5 clusters was performed. Finally, a trend analysis was elaborated to evaluate the past and current course of research on cafeteria diet and behavior variables.

## Results

### 0.5 Performance analysis

Table S1 lists the main results of the performance analysis. As a result of the analysis a total of 85 original articles from 2013 to 2023 with an annual growth rate of 1.34 % and 18.18 average citations per article, with an average citation per year of 2.8. 457 authors with 6.52 co-authors per article were retrieved, and the international co-authorship rate was 12.94%. No single-authored articles were reported. The keywords provided by the authors were 423 and 677 keywords were provided by the journal source. Three were the databases accessed, one open source [Pubmed (31)] and two reached by institutional subscription: Web of Science (32) and Scopus (33).

### 0.6 Citation-related metrics

Citations can assess the importance of scientific performance, the number of citations of scientist’s publications has been shown to correlate with other assessments of the impact or influence of scientists, such as prizes, awards, and distinctions (34). In this analysis articles between 0 to 115 citations were included, with a cumulative sum of 1545 and a mean of 18.17. From 2013 to 2023, the most productive year was 2018 with 10 articles, and the least productive year was 2020 with 5 articles. All 85 articles are presented in Table S2 with information on the title, journal, year of publication, citations, authors, and database from which they were extracted. Table 2 shows the top 10 articles with the highest number of citations.

**Table 2.**
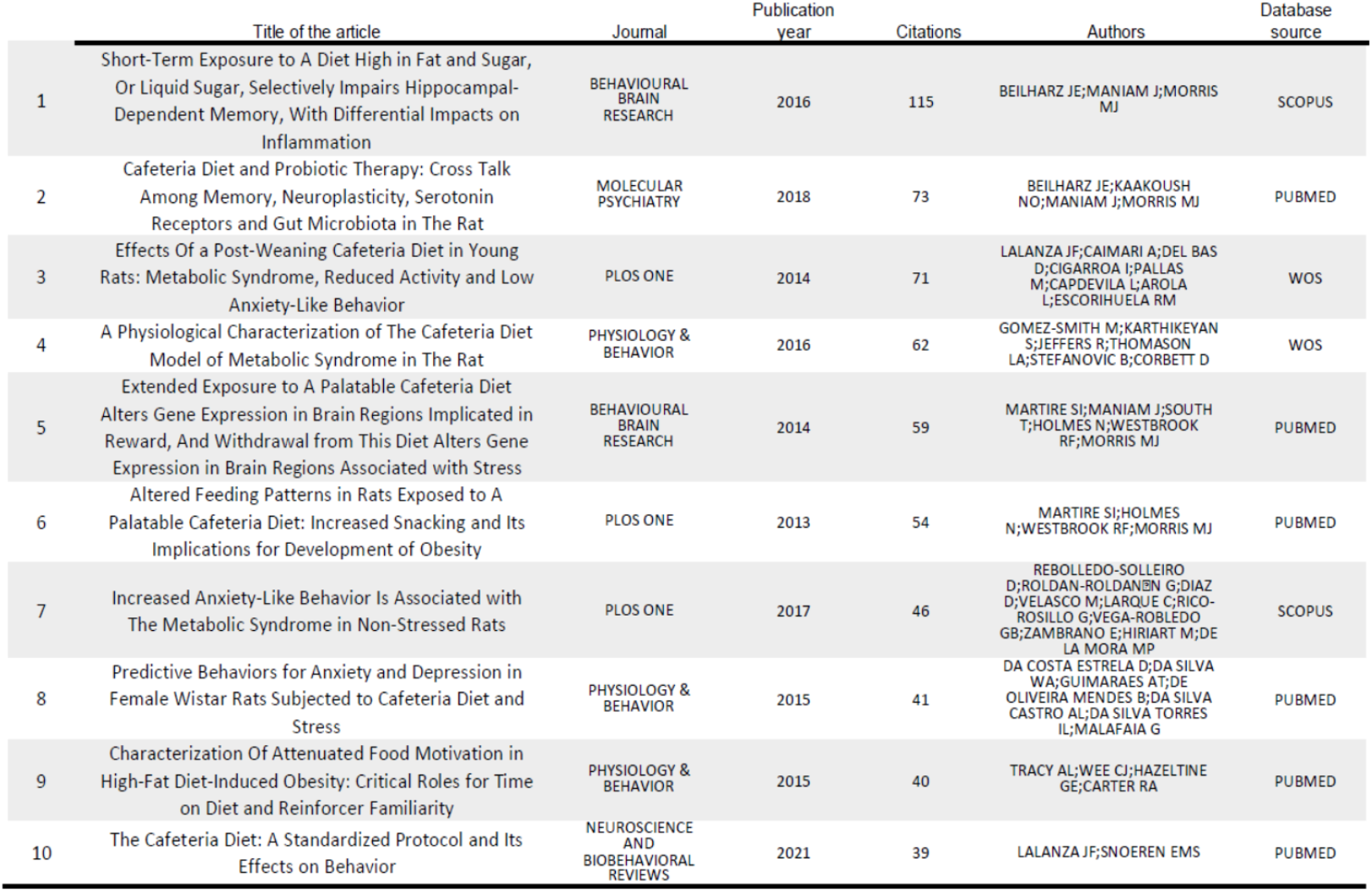
10 most cited articles. Journal, publication year, number of citations until 2023, authors and database source are enlisted.

### 0.7 Citation and publication-related metrics

Of the 457 total authors, only 15 had an h-index higher than 3 (Table 3). Morris and Westbrook had the highest h-index, g-index, and number of articles, with 447 and 227 citations, respectively. The m index varied from 0.90 to 0.30.

**Table 3.**
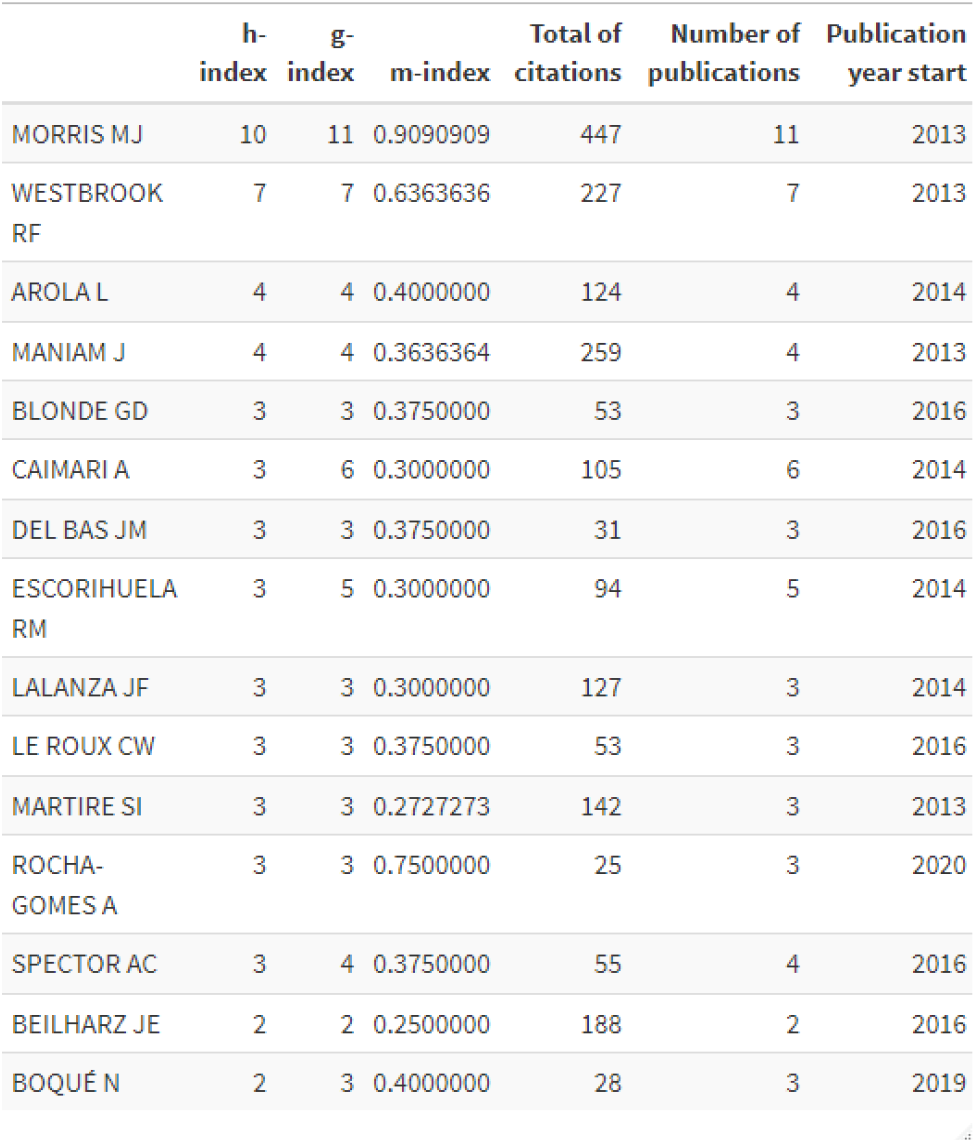
15 most productive authors. H-index, g-index, m-index, and total citations for the number of publications since the beginning of the year of publication of the 15 most productive authors with an h-index ¿3 are enlisted.

|  | h-index | g-index | m-index | Total of citations | Number of publications | Publication year start |
| --- | --- | --- | --- | --- | --- | --- |
| MORRIS MJ | 10 | 11 | 0.9090909 | 447 | 11 | 2013 |
| WESTBROOK RF | 7 | 7 | 0.6363636 | 227 | 7 | 2013 |
| AROLA L | 4 | 4 | 0.4000000 | 124 | 4 | 2014 |
| MANIAM J | 4 | 4 | 0.3636364 | 259 | 4 | 2013 |
| BLONDE GD | 3 | 3 | 0.3750000 | 53 | 3 | 2016 |
| CAIMARI A | 3 | 6 | 0.3000000 | 105 | 6 | 2014 |
| DEL BAS JM | 3 | 3 | 0.3750000 | 31 | 3 | 2016 |
| ESCORIHUELA RM | 3 | 5 | 0.3000000 | 94 | 5 | 2014 |
| LALANZA JF | 3 | 3 | 0.3000000 | 127 | 3 | 2014 |
| LE ROUX CW | 3 | 3 | 0.3750000 | 53 | 3 | 2016 |
| MARTIRE SI | 3 | 3 | 0.2727273 | 142 | 3 | 2013 |
| ROCHA-GOMES A | 3 | 3 | 0.7500000 | 25 | 3 | 2020 |
| SPECTOR AC | 3 | 4 | 0.3750000 | 55 | 4 | 2016 |
| BEILHARZ JE | 2 | 2 | 0.2500000 | 188 | 2 | 2016 |
| BOQUÉ N | 2 | 3 | 0.4000000 | 28 | 3 | 2019 |

The top 25 authors with the highest number of articles published about cafeteria diet and behavior were grouped by country, resulting in 5 countries. Spain had 40% (10 authors), followed by Brazil with 24% (6 authors) and Australia with 20% (5 authors) of the top 25 authors (Fig 2). The number of articles in which the author has participated is shown in parenthesis and is also differentiated by the height of the bars.

**Fig 2.**
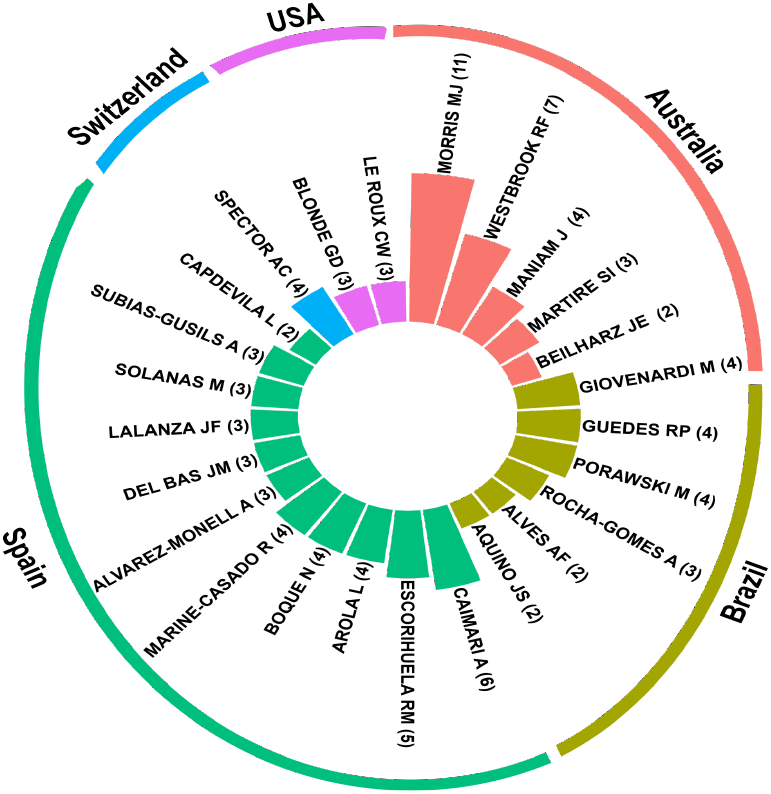
Most productive 25 authors. The circle bar chart shows the 25 most productive authors grouped by country: Australia (orange), Brazil (olive green), Spain (green), Switzerland (blue), and USA (pink). Authors are also ordered by highest to lowest number of articles published within countries.

### 0.8 Science mapping

#### 0.8.1 Citation analysis

A total of 85 articles were produced by 20 countries (Fig 3a), with 22 articles from Brazil that correspond to 26% of the total, followed by Australia and Spain with 13 and 12 articles, respectively. From 2013 to 2023, the most cited papers were produced by Australia with 173 citations as is shown in Fig 3b.

**Fig 3.**
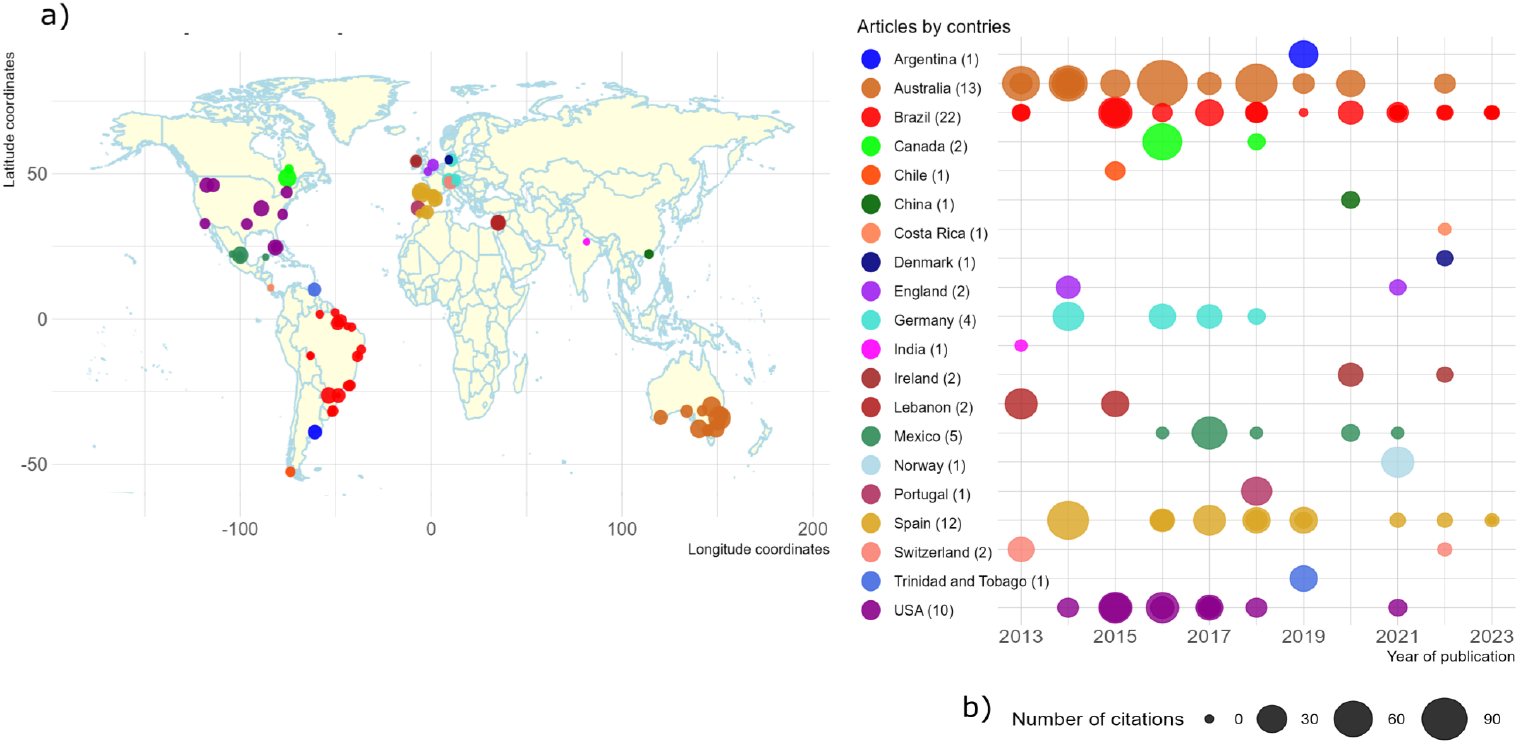
Citation country analysis. a) Article production per country and number of citations is plotted in dots. Bigger circles indicate more citations, as shown in b). b) Number of articles and citations per country and year.

Of 56 institutions of corresponding author affiliations, only 14 institutions, representing 17% of the total, published 2 articles. The University of New South Wales, Universitat Autonoma de Barcelona and the Universidade Federal de Ciências da Saúde de Porto Alegre are the most productive institutions with 10, 5 and 4 articles, respectively.

Although the impact factor is a metric sought after by academics, it is a dynamic measure that changes. Thus, to answer whether the impact factor of the journals is related to the number of citations obtained up to the time of data collection for this study, a Pearson’s Correlation (r 0.095) was applied, suggesting a weak but positive relationship between the IF and citations. Therefore, we applied a Simple Linear Regression Model (Figure 5a) for the IF and number of citations as dependent and independent variables. The Intercept was 3.58 (SD 0.25 p ¡ 0.001) but no significance for the number of citations was found (p = 0.385 R^2^ = 0.009 83 DOF). Thus, no positive linear relationship between IF and the number of citations was found. Of the total 49 journals, only 12 journals (24.5%) received 2 articles.

**Fig 4.**
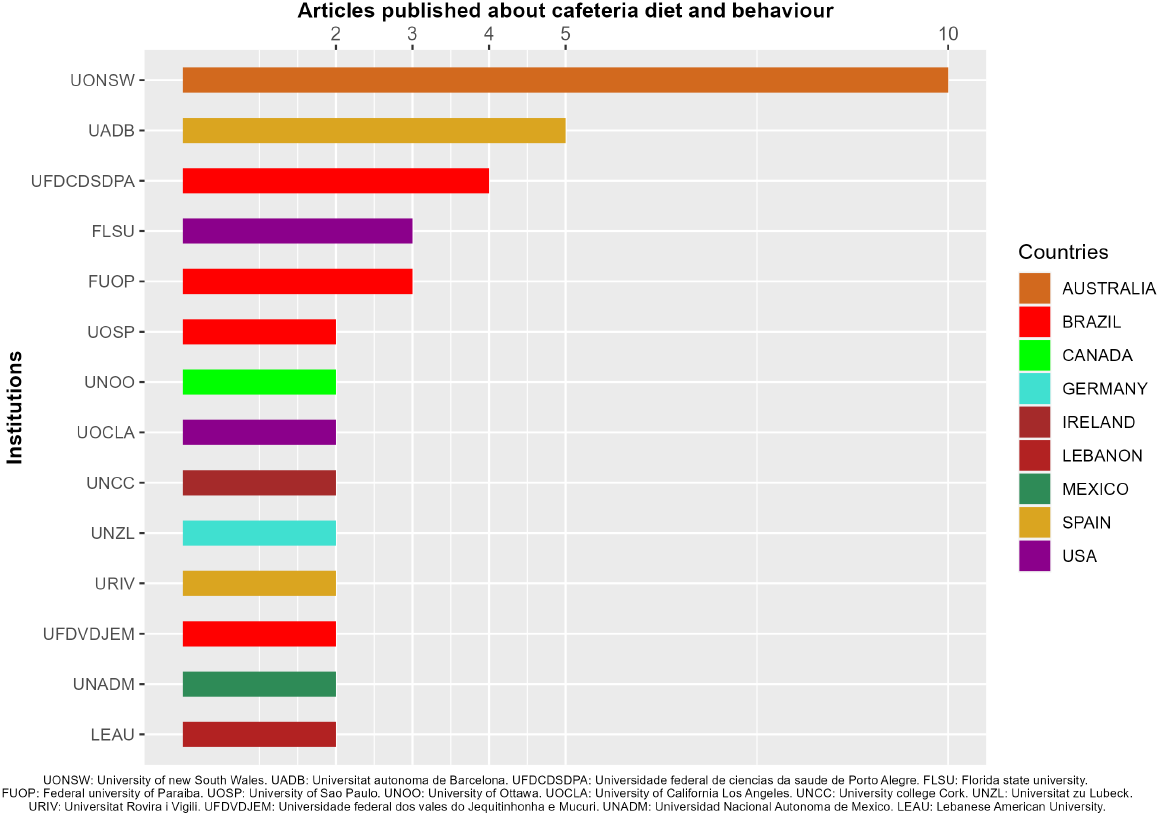
Articles published by Institutions. 14 institutions with at least 2 published articles, colored by country.

**Fig 5.**
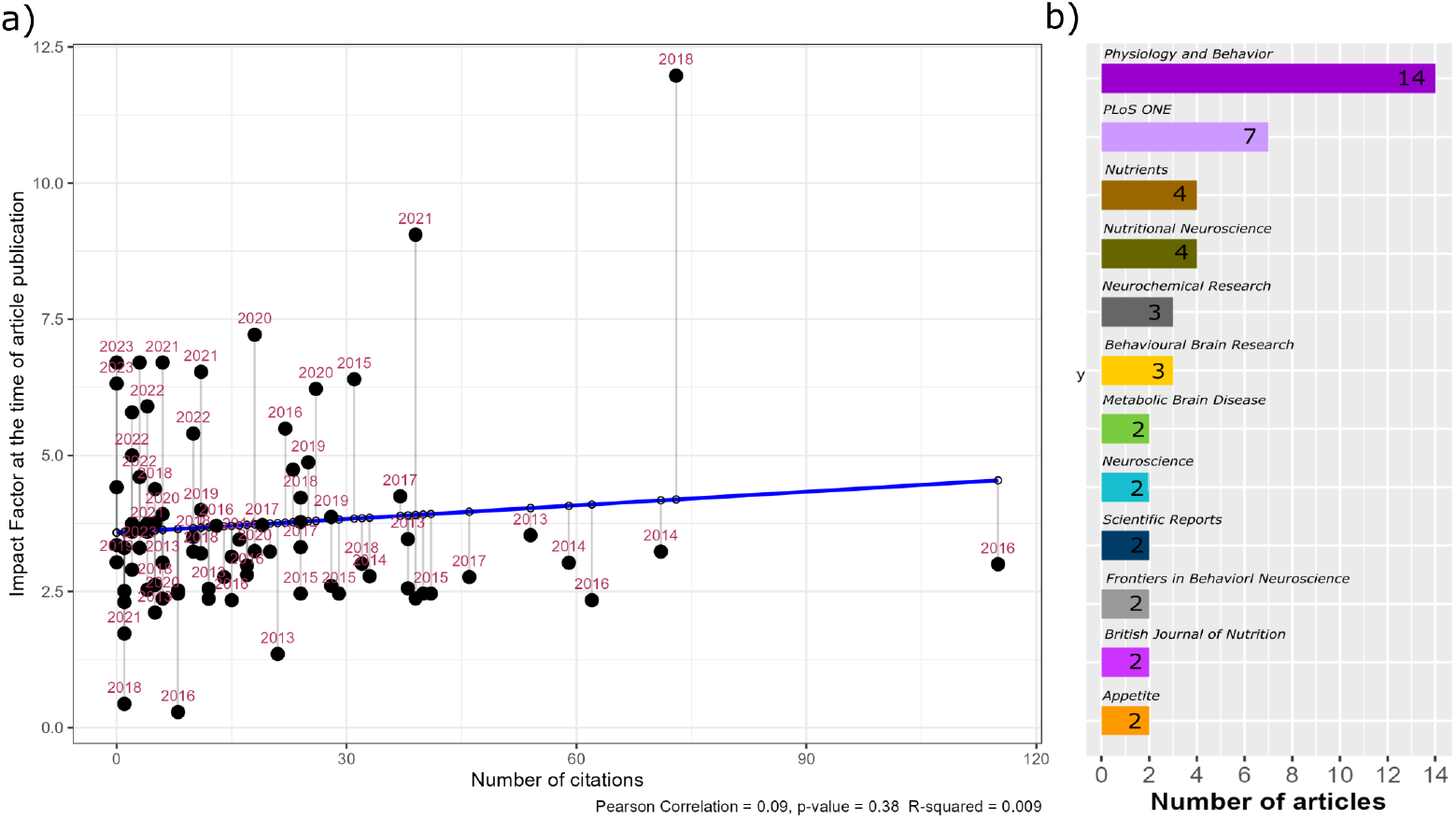
Simple Linear Regression Model (SLRM) of the IF and number of citations. a) The SLRM was adjusted to determine the relationship between the IF and citations (p = 0.38, R2(0.09). No linear relationship between the IF and citations was found. Year of publication per article is shown. b) Top submitting journals, with 2 articles published. The number in each bar indicates the total of articles published per journal.

#### 0.8.2 Co-word analysis

A total of 57 terms were extracted and weighted to perform the document term-matrix (dfm). The strength of each term is enlisted in (Table S3). A map of the distance and node location is graphed in Fig S1. The MCA method was composed of two dimensions that described the 44.1% and 25.05% of the explained variance. A total of five clusters were found.The lilac cluster consists of the terms: nucleus accumbens, memory and high-fat diet. The blue cluster includes dietary fats, food preference, high-fat, energy intake, sucrose, adipose tissue, obesity, female, male, stress, leptin, gene expression, anxiety, and inflammation. The green cluster groups: receptor, dopamine, cognition, drug effects, pharmacology, administration & dosage, etiology, physiology, animal, metabolism and again the terms genetics, adiposity, diet and feeding behavior. The mustard cluster is composed of weight gain, body weight, disease patterns, brain-derived neurotrophic factor, blood, and hippocampus. The red cluster includes insulin, corticosterone, calorie intake, blood glucose, glucose level, adult, article, and controlled study.

Finally, a bi-factor analysis allowed us to model the trending topics (35). This model is used to analyze the relationships between observed variables and underlying latent factors, in this case, trends over 10 years. Anxiety is the trending topic related to cafeteria diet and behavior since 2017 to this date. Its trend appears constant but has a low frequency of appearance. On the other hand, metabolism had the highest frequency term, but its trend line only remained from 2017 to 2019. High-fat and sucrose were terms with long trend lines, from 2013 to 2019 and from 2015 to 2021, respectively.

**Fig 6.**
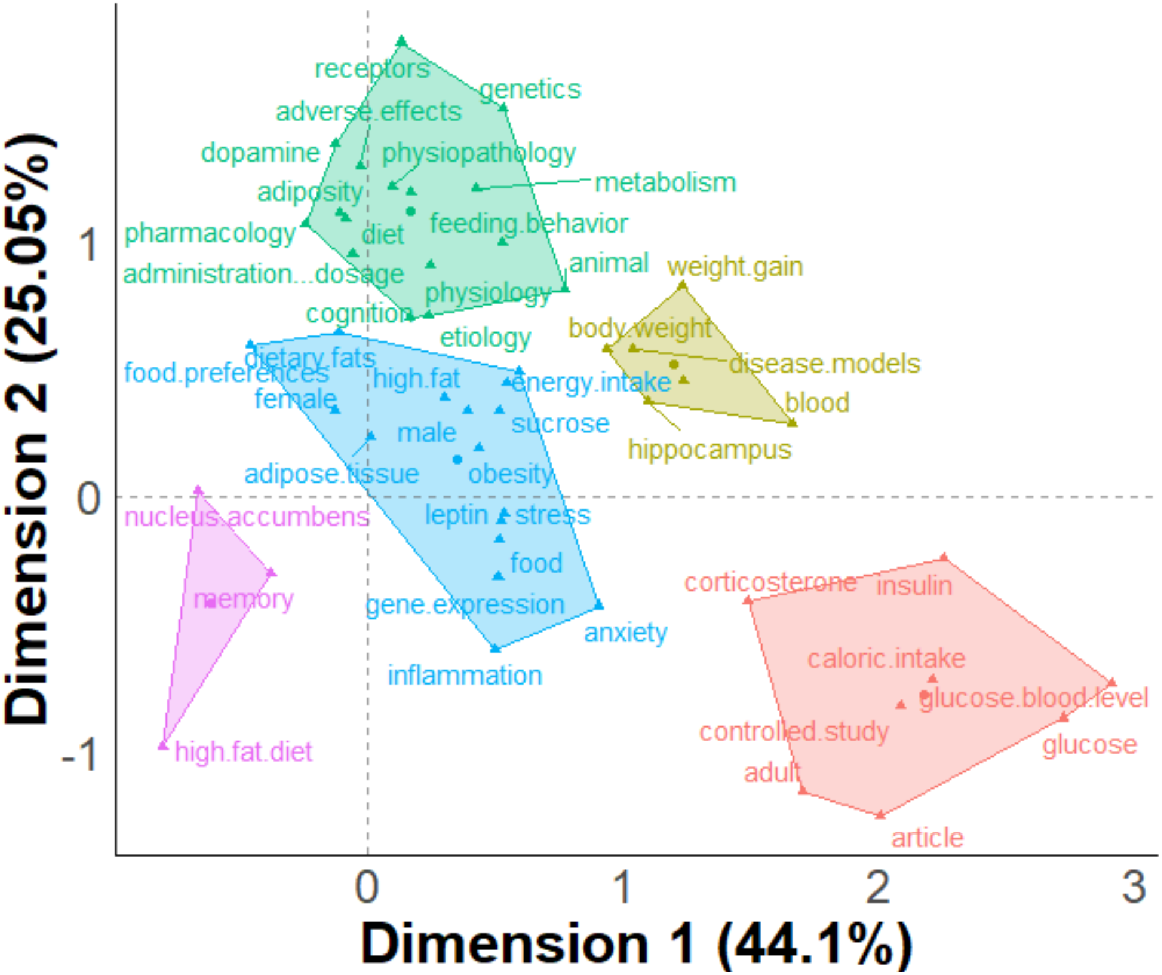
Conceptual structure of the articles by multiple correspondence analysis (MCA). The strength of the dfm was modeled with a conceptual structure map, with an MCA approach with 5 k-means.

**Fig 7.**
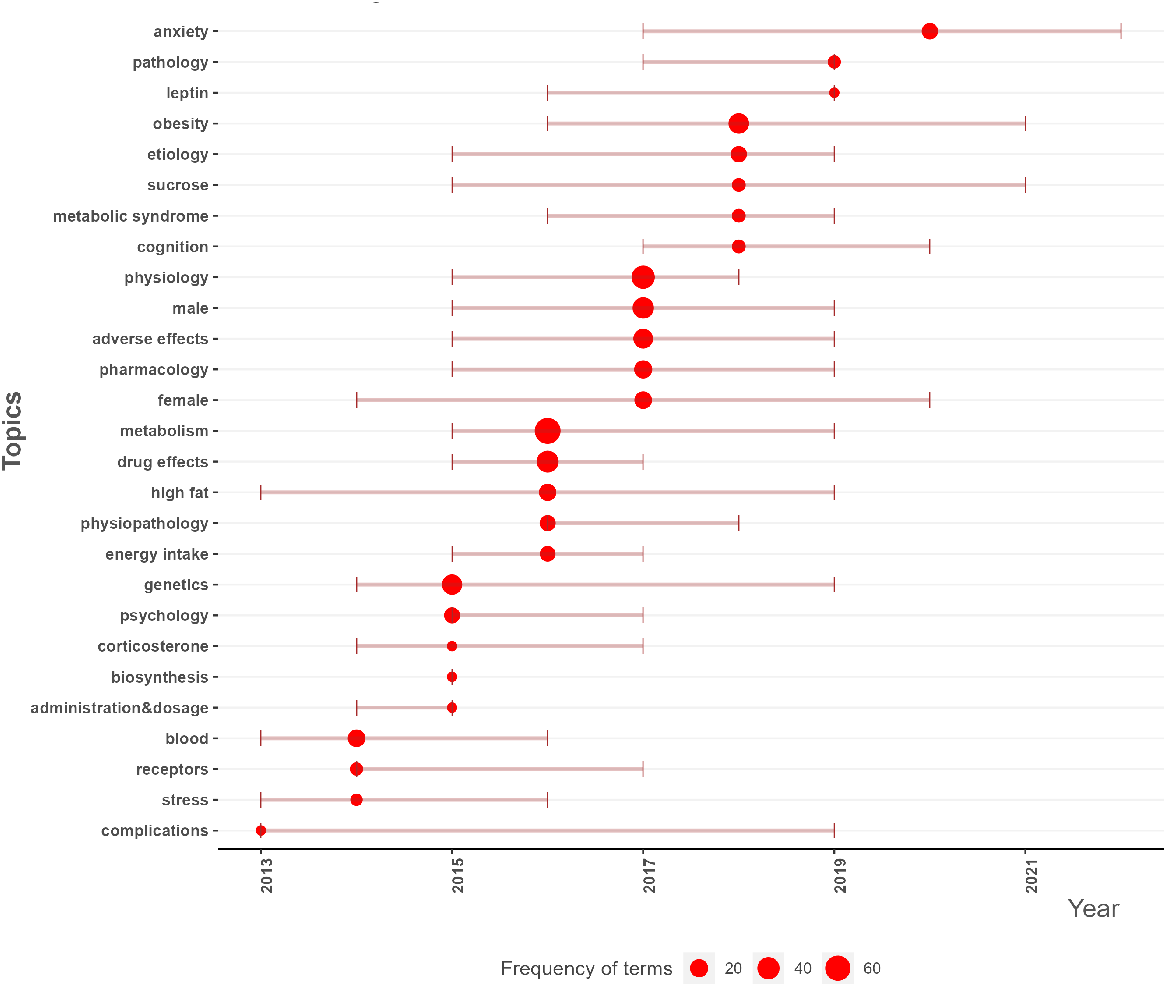
Trending topics. Topics and highest frequency of terms in article database from the last 10 years. The size of the circles indicates the frequency of terms and the length of the lines shows the presence of the topic over the years.

## Discussion

Bibliometric analysis is a rigorous method that allows to unravel the intellectual structure of a research field. It also sheds light on the performance of authors, institutions, and countries, while uncovering emerging trends and the relationship between them (22). This study performed a bibliometric analysis on the rising field of the impact of CAF diet on metabolism, neurobiology and behavior. Considering that there is a disparity of information among several databases (36), three databases were consulted: Pubmed, Scopus and Web of Science. Of the initial 365 articles, 85 met the eligibility criteria, publication and citation metrics were captured and processed for top authors, journals, institutions and countries. Then, we explored the relationship between terms and performed science mapping to identify trending topics. To our knowledge, this is the first bibliometric analysis performed in this research field. This study will help the scientific community to access the information generated over the last 10 years in this research field, and to consider future directions. It will also help to optimize the time spent in research, which has decreased over time (37). The 85 articles are listed in Table S1 ordered from highest to lowest in citation, and as expected the top articles were led by the top corresponding authors, and the journals with the highest IF (34). Beilharz et al., (2016) was the most cited article in this study (115 citations), where a detailed exploration on the impact of CAF diet on metabolism, neurobiology and behavior was conducted. The second most cited was Beilharz et al.,(2018), which links the aforementioned disturbances with one of the trend topics in research of obesity etiology: microbiota as a target for therapeutic approaches (38). The corresponding author of the two mentioned articles is Morris M.J., who leads the author analysis by being the author with the highest h-index (29). This index, as reported in Table 3, does not depend on the years of scientific production. Authors such as Boque, N. with a publication start in 2019, 31 citations and 4 articles, reached an h-index similar to authors who have more years or more accumulated citations. Third most cited publication (71 citations) is also published by a group of highly prolific and cited authors: Lalanza et al. (2014). The most productive countries (Australia, Brazil, and Spain) are home of the most productive and cited institutions (UONSW, UADB, UFDCDSDPA). Accordingly, Australia (30.57%), Brazil (19.24%), and Spain (25.49%). Journal IF has been considered directly correlated to higher citations (42), although the impact factor is dynamic, it is considered an adequate estimator to predict future citation success (34). However, we did not find a positive correlation between impact factor and the number of citations. Other groups consider the use of highly dynamic journal IF as inadequate and have proposed other metrics to complement journal IF or to predict publication long-term impact (43);(44);(45). According to these results of the trend analysis, the keywords pharmacology, drug effects, cognition and metabolism are trending topics that increased in frequency 2015-2017. In MCA and trend analysis, anxiety is one of the keywords, most commonly used from 2017 to date. Finally, it is important to mention that this study has several limitations. Only original articles in English were included. In addition, by including just 10 years of results, it is to be expected that the list of the most cited publications will change over time, as a consequence of the exponential expansion of research. Lastly, the number of citations is only a proxy indicator of scientific impact and may be affected by other factors, such as accessibility and journal reputations.

## Conclusion

The present study explores the performance of authors, countries, and institutions, and performs a science mapping to carry out a multiple correspondence analysis and a trend analysis of the research field. The most cited articles and authors are described, which may guide those interested in considering the evolution of the research field over the last 10 years. It will be complemented by information provided by the trend analysis that may reinforce decisions to explore missing relationships to complement multifactorial considerations of obesity and metabolic alterations of the cafeteria diet.

## Acknowledgments

A Lopez-Castro is a doctoral student from the Programa de Doctorado en Ciencias Biomédicas, Universidad Nacional Autónoma de México (UNAM) and has received CONAHCYT fellowship 1003251.

