## supplementary for "A bibliometric analysis of neurobiological and behavioral disturbances of cafeteria diet interventions"

### SUPPLEMENTARY MATERIAL

| | Title | Journal | Year | Cites | Authors | DB |
|---|---|---|---|---|---|---|
| 1 | Short-term exposure to a diet high in fat and sugar, or liquid sugar, selectively impairs hippocampal-dependent memory, with differential impacts on inflammation | Behavioural Brain Research | 2016 | 115 | Beilharz Je;Maniam J;Morris Mj | SCOPUS |
| 2 | Cafeteria diet and probiotic therapy: cross talk among memory, neuroplasticity, serotonin receptors and gut microbiota in the rat | Molecular Psychiatry | 2018 | 73 | Beilharz Je;Kaakoush No;Maniam J;Morris Mj | PUBMED |
| 3 | Effects of a post-weaning cafeteria diet in young rats: metabolic syndrome, reduced activity and low anxiety-like behaviour | Plos One | 2014 | 71 | Lalanza Jf;Caimari A;Del Bas D;Cigarroa I;Pallas M;Capdevila L;Arola L;Escorihuela Rm | WOS |
| 4 | A physiological characterization of the cafeteria diet model of metabolic syndrome in the rat | Physiology & Behavior | 2016 | 62 | Gomez-Smith M;Karthikeyan S;Jeffers R;Thomason La;Stefanovic B;Corbett D | WOS |
| 5 | Extended exposure to a palatable cafeteria diet alters gene expression in brain regions implicated in reward, and withdrawal from this diet alters gene expression in brain regions associated with stress | Behavioural Brain Research | 2014 | 59 | Martire Si;Maniam J;South T;Holmes N;Westbrook Rf;Morris Mj | PUBMED |
| 6 | Altered feeding patterns in rats exposed to a palatable cafeteria diet: increased snacking and its implications for development of obesity | Plos One | 2013 | 54 | Martire Si;Holmes N;Westbrook Rf;Morris Mj | PUBMED |
| 7 | Increased anxiety-like behavior is associated with the metabolic syndrome in non-stressed rats | Plos One | 2017 | 46 | Rebolledo-Solleiro D;Roldan-RoldanN G;Diaz D;Velasco M;Larque C;Rico-Rosillo G;Vega-Robledo Gb;Zambrano E;Hiriart M;De La Mora Mp | SCOPUS |
| 8 | Predictive behaviors for anxiety and depression in female wistar rats subjected to cafeteria diet and stress | Physiology & Behavior | 2015 | 41 | Da Costa Estrela D;Da Silva Wa;Guimaraes At;De Oliveira Mendes B;Da Silva Castro Al;Da Silva Torres Il;Malafaia G | PUBMED |
| 9 | Characterization of attenuated food motivation in high-fat diet-induced obesity: critical roles for time on diet and reinforcer familiarity | Physiology & Behavior | 2015 | 40 | Tracy Al;Wee Cj;Hazeltine Ge;Carter Ra | PUBMED |
| 10 | The cafeteria diet: a standardized protocol and its effects on behavior | Neuroscience And Biobehavioral Reviews | 2021 | 39 | Lalanza Jf;Snoeren Ems | PUBMED |
| 11 | Roux-en-y gastric bypass in rats progressively decreases the proportion of fat calories selected from a palatable cafeteria diet | American Journal Of Physiology. Regulatory, Integrative And Comparative Physiology | 2016 | 39 | Mathes Cm;Letourneau C;Blonde Gd;Le Roux Cw;Spector Ac | PUBMED |
| 12 | Cafeteria diet impairs expression of sensory-specific satiety and stimulus-outcome learning | Frontiers In Psychology | 2014 | 38 | Reichelt Ac;Morris Mj;Westbrook Rf | WOS |
| 13 | A cafeteria diet modifies the response to chronic variable stress in rats | Stress (Amsterdam, Netherlands) | 2013 | 38 | Zeeni N;Daher C;Fromentin G;Tome D;Darcel N;Chaumontet C | PUBMED |
| 14 | Effects of bingeing on fat during adolescence on the reinforcing effects of cocaine in adult male mice | Neuropharmacology | 2017 | 37 | Carmen Blanco-Gandia M;Cantacorps L;Aracil-Fernandez A;Montagud-Romero S;Aguilar J;Valverde O;Minarro M | WOS |
| 15 | The impact of cafeteria diet feeding on physiology and anxiety-related behaviour in male and female sprague-dawley rats of different ages | Pharmacology, Biochemistry, And Behavior | 2014 | 33 | Warneke W;Klaus S;Fink H;Langley-Evans Sc;Voigt Jp | PUBMED |

| | | | | | | |
|---|---|---|---|---|---|---|
| **16** | Cafeteria-diet effects on cognitive functions, anxiety, fear response and neurogenesis in the juvenile rat | Neurobiology Of Learning and Memory | 2018 | 32 | Ferreira A;Castro Jp;Andrade Jp;Dulce Madeira M;Cardoso A | SCOPUS |
| **17** | Involvement of endogenous enkephalins and ï’-endorphin in feeding and diet-induced obesity | Neuropsychopharmacology: Official Publication Of The American College Of Neuropsychopharmacology | 2015 | 31 | Mendez Ia;Ostlund Sb;Maidment Nt;Murphy Np | PUBMED |
| **18** | Effects of long-term cycling between palatable cafeteria diet and regular chow on intake, eating patterns, and response to saccharin and sucrose | Physiology & Behavior | 2015 | 29 | Martire Si;Westbrook Rf;Morris Mj | PUBMED |
| **19** | Cafeteria diet induces progressive changes in hypothalamic mechanisms involved in food intake control at different feeding periods in female rats | Molecular And Cellular Endocrinology | 2019 | 28 | Lazzarino Gp;Acutain Mf;Canesini G;Andreoli Mf;Ramos Jg | PUBMED |
| **20** | Effects of palatable cafeteria diet on cognitive and noncognitive behaviors and brain neurotrophins' levels in mice | Metabolic Brain Disease | 2015 | 28 | Leffa Dd;Valvassori Ss;Varela Rb;Lopes-Borges J;Daumann F;Longaretti Lm;Dajori Al;Quevedo J;Andrade Vm | PUBMED |
| **21** | Intermittent cafeteria diet identifies fecal microbiome changes as a predictor of spatial recognition memory impairment in female rats | Translational Psychiatry | 2020 | 26 | Leigh Sj;Kaakoush No;Bertoldo Mj;Westbrook Rf;Morris Mj | PUBMED |
| **22** | Impact of different hypercaloric diets on obesity features in rats: a metagenomics and metabolomics integrative approach | The Journal Of Nutritional Biochemistry | 2019 | 25 | Gual-Grau A;Guirro M;Mayneris-Perxachs J;Arola L;Boque N | PUBMED |
| **23** | Cafeteria diet induces neuroplastic modifications in the nucleus accumbens mediated by microglia activation | Addiction Biology | 2018 | 24 | Gutierrez-Martos M;Girard B;Mendonca-Netto S;Perroy J;Valjent E;Maldonado M | WOS |
| **24** | Environmental enrichment and cafeteria diet attenuate the response to chronic variable stress in rats | Physiology & Behavior | 2015 | 24 | Zeeni N;Bassil M;Fromentin G;Chaumontet C;Darcel N;Tome D;Daher Cf | SCOPUS |
| **25** | Effects of cafeteria diet and high fat diet intake on anxiety, learning and memory in adult male rats | Nutritional Neuroscience | 2017 | 24 | Pini Rtb;Ferreira Do Vales Ldm;Braga Costa Tm;Almeida Ss | PUBMED |
| **26** | Cafeteria-diet induced obesity results in impaired cognitive functioning in a rodent model | Heliyon | 2019 | 24 | Lewis Ar;Singh S;Youssef Ff | PUBMED |
| **27** | Effects of a cafeteria diet on delay discounting in adolescent and adult rats: alterations on dopaminergic sensitivity | Journal Of Psychopharmacology (Oxford, England) | 2017 | 23 | Robertson Sh;Rasmussen Eb | PUBMED |
| **28** | The brain renin-angiotensin system plays a crucial role in regulating body weight in diet-induced obesity in rats | British Journal Of Pharmacology | 2016 | 22 | Winkler M;Schuchard J;Stoelting I;Vogt Fm;Barkhausen J;Thorns C;Bader M;Raasch W | WOS |
| **29** | The selective orexin receptor 1 antagonist act-335827 in a rat model of diet-induced obesity associated with metabolic syndrome | Frontiers In Pharmacology | 2013 | 21 | Steiner Ma;Sciarretta C;Pasquali A;Jenck F | WOS |
| **30** | Rats eat a cafeteria-style diet to excess but eat smaller amounts and less frequently when tested with chow | Plos One | 2014 | 20 | South T;Holmes Nm;Martire Rf;Morris Mj | WOS |
| **31** | Early effects of a high-caloric diet and physical exercise on brain volumetry and behavior: a combined mri and histology study in mice | Brain Imaging And Behavior | 2017 | 19 | Sack M;Lenz Jn;Jakovcevski M;Biedermann Sv;Falfan-Melgoza C;Deussing J;Bielohuby M;Bidlingmaier M;Pfister F;Stalla Gk;Sartorius A;Gass P;Weber-Fahr W;Fuss J;Auer Mk | WOS |

| | | | | | | |
|---|---|---|---|---|---|---|
| **32** | Adolescent dietary manipulations differentially affect gut microbiota composition and amygdala neuroimmune gene expression in male mice in adulthood | Brain Behavior And Immunity | 2020 | 18 | Fulling C;Lach G;Bastiaanssen F;O'donovan An;Ventura-Silva C;Dinan Tg;Cryan Jf | WOS |
| **33** | Cafeteria diet administered from lactation to adulthood promotes a change in risperidone sensitivity on anxiety, locomotion, memory, and social interaction of wistar rats | Physiology & Behavior | 2020 | 18 | Teixeira Ae;Rocha-Gomes A;Pereira Dos Santos T;Amaral Bls;Da Silva Aa;Malagutti Ar;Leite Frf;Stuckert-Seixas Sr;Riul Tr | PUBMED |
| **34** | Treadmill intervention attenuates the cafeteria diet-induced impairment of stress-coping strategies in young adult female rats | Plos One | 2016 | 17 | Cigarroa I;Lalanza Jf;Caimari A;Del Bas Jm;Capdevila L;Arola L;Escorihuela Rm | PUBMED |
| **35** | Naloxone treatment alters gene expression in the mesolimbic reward system in 'junk food' exposed offspring in a sex-specific manner but does not affect food preferences in adulthood | Physiology & Behavior | 2014 | 17 | Gugusheff Jr;Ong Zy;Muhlhausler Bs | PUBMED |
| **36** | Impact of cafeteria feeding during lactation in the rat on novel object discrimination in the offspring | British Journal Of Nutrition | 2014 | 16 | Wright Tm;King Mv;Davey Sc;Voigt Jpw | WOS |
| **37** | Modern 'junk food' and minimally-processed 'natural food' cafeteria diets alter the response to sweet taste but do not impair flavor-nutrient learning in rats | Physiology & Behavior | 2016 | 15 | Palframan Km;Myers Kp | PUBMED |
| **38** | Western diet chow consumption in rats induces striatal neuronal activation while reducing dopamine levels without affecting spatial memory in the radial arm maze | Frontiers In Behavioral Neuroscience | 2017 | 15 | Nguyen Jc;Ali Sf;Kosari S;Woodman Ol;Spencer Sj;Killcross As;Jenkins Ta | PUBMED |
| **39** | Junk food diet-induced obesity increases d2 receptor autoinhibition in the ventral tegmental area and reduces ethanol drinking | Plos One | 2017 | 14 | Cook Jb;Hendrickson Lm;Garwood Gm;Toungate Km;Nania Cv;Morikawa H | PUBMED |
| **40** | The intake of high-fat diets induces an obesogenic-like gene expression profile in peripheral blood mononuclear cells, which is reverted by dieting | The British Journal Of Nutrition | 2016 | 13 | Reynes B;GarcãA-Ruiz E;Palou A;Oliver P | PUBMED |
| **41** | Early life stress and post-weaning high fat diet alter tyrosine hydroxylase regulation and at1 receptor expression in the adrenal gland in a sex dependent manner | Neurochemical Research | 2013 | 12 | Bobrovskaya L;Maniam J;Ong Lk;Dunkley Pr;Morris Mj | PUBMED |
| **42** | Cafeteria diet during the gestation period programs developmental and behavioral courses in the offspring | International Journal Of Developmental Neuroscience : The Official Journal Of The International Society For Developmental Neuroscience | 2018 | 12 | Ribeiro Acaf;Batista Th;Veronesi Vb;Giusti-Paiva A;Vilela Fc | PUBMED |
| **43** | Potentially obesogenic diets alter metabolic and neurobehavioural parameters in wistar rats: a comparison between two dietary models | Journal Of Affective Disorders | 2021 | 11 | Bonfim Thf;Tavares Rl;De Vasconcelos Mha;Gouveia M;Nunes Pc;Soares Nl;Alves Rc;De Carvalho Jlp;Alves Af;Pereira Ra;Cardoso Ga;Silva As;Aquino Js | SCOPUS |
| **44** | Intake of an obesogenic cafeteria diet affects body weight, feeding behavior, and glucose and lipid metabolism in a photoperiod-dependent manner in f344 rats | Frontiers In Physiology | 2018 | 11 | Marine-Casado R;Domenech-Coca C;Del Bas Jm;Blade C;Arola L;Caimari A | WOS |
| **45** | Effects of cafeteria diet on memory and hippocampal oxidative stress in a rat model of alzheimer-like disease: neuroprotection of green tea supplementation | Journal Of Functional Foods | 2018 | 11 | Schimidt Hl;Garcia A;Martins M;Soares Mb;Cibin Pb;Carpes Fp | WOS |
| **46** | Pattern of access to cafeteria-style diet determines fat mass and degree of spatial memory impairments in rats | Scientific Reports | 2019 | 11 | Kendig Md;Westbrook Rf;Morris Mj | PUBMED |

| | | | | | | |
|---|---|---|---|---|---|---|
| 47 | Effect of cafeteria diet history on cue-, pellet-priming-, and stress-induced reinstatement of food seeking in female rats | Plos One | 2014 | 10 | Chen Yw;Fiscella Ka;Bacharach Dj | WOS |
| 48 | Pattern of access determines influence of junk food diet on cue sensitivity and palatability | Appetite | 2018 | 10 | Kosheleff Ar;Araki J;Hsueh J;Le A;Quizon K;Ostlund Sb;Maidment Np | WOS |
| 49 | Chronic exposure to cafeteria-style diet in rats alters sweet taste preference and reduces motivation for, but not 'liking' of sucrose | Appetite | 2022 | 10 | Fam J;Clemens Kj;Westbrook Rf;Morris Mj;Kendig Md | PUBMED |
| 50 | Behavioral characterization of a model of differential susceptibility to obesity induced by standard and personalized cafeteria diet feeding | Physiology & Behavior | 2015 | 8 | Gac L;Kanaly V;Ramirez V;Teske Ja;Pinto Mp;Perez-Leighton Ce | PUBMED |
| 51 | Liraglutide suppression of caloric intake competes with the intake-promoting effects of a palatable cafeteria diet, but does not impact food or macronutrient selection | Physiology & Behavior | 2017 | 8 | Hyde Km;Blonde Gd;Le Roux Cw;Spector Ac | PUBMED |
| 52 | Renal inflammatory and oxidative and metabolic changes after 6 weeks of cafeteria diet in rats | Jornal Brasileiro De Nefrologia | 2016 | 8 | Navarro Me;Santos Kc;Nascimento Af;Francisqueti Fv;Minatel Io;Pierine Dt;Luvizotto Ra;Ferreira Al;Campos Dh;Correa Cr | PUBMED |
| 53 | Aqueous extract of pomegranate enriched in ellagitannins prevents anxiety-like behavior and metabolic changes induced by cafeteria diet in an animal model of menopause | Neurochemistry International | 2020 | 6 | Estrada-Camarena Em;Lopez-Rubalcava C;Ramirez-Rodriguez Gb;Pulido D;Cervantes-Anaya N;Azpilcueta-Morales G;Granados-Juarez A;Vega-Rivera Nm;Islas-Preciado D;Trevino S;De Gortari P;Gonzalez-Trujano C | WOS |
| 54 | Meal patterns and food choices of female rats fed a cafeteria-style diet are altered by gastric bypass surgery | Nutrients | 2021 | 6 | Blonde Gd;Price Rk;Le Roux Cw;Spector Ac | PUBMED |
| 55 | Prevention of metabolic disorders and reproductive performance deficits by the blockade of angiotensin ii at1 receptor in female rats fed with cafeteria diet | Physiology & Behavior | 2013 | 6 | Sagae Sc;Lubaczeuski C;Zacharias P;Bonfleur Ml;Franci Cr;Sanvitto Gl | PUBMED |
| 56 | Cafeteria-style feeding trials provide new insights into the diet and nutritional strategies of the black snub-nosed monkey (rhinopithecus strykeri): implications for conservation | American Journal Of Primatology | 2020 | 6 | Yang Y;Li Q;Garber Pa;Grueter Cc;Ren G;Wang X;Huang Z;Xiang Z;Xiao W;Behie A | PUBMED |
| 57 | Palatable food dampens the long-term behavioral and endocrine effects of juvenile stressor exposure but may also provoke metabolic syndrome in rats | Frontiers In Behavioral Neuroscience | 2018 | 5 | Ali Ef;Mackay Jc;Graitson S;James Js;Cayer C;Audet Mc;Kent P;Abizaid A;Merali Z | WOS |
| 58 | Influence of at1 blockers on obesity and stress-induced eating of cafeteria diet | Journal Of Endocrinology | 2018 | 5 | Gustaityte V;Winkler M;Stolting I;Raasch W | SCOPUS |
| 59 | Hypercaloric high-lipid diet and brain development: effects on cortical spreading depression in adult rats | Nutritional Neuroscience | 2013 | 5 | Da Silva Germano Pc;De Lima E Silva D;Soares Gde S;Dos Santos Aa;Guedes Rc | PUBMED |
| 60 | Proanthocyanidins limit adipose accrual induced by a cafeteria diet, several weeks after the end of the treatment | Genes | 2019 | 5 | Gines I;Gil-Cardoso K;Serrano J;Casanova-Marti A;Lobato M;Terra X;Blay Mt;Ardevol A;Pinent M | PUBMED |
| 61 | High-fat diet and fructose drink introduced after weaning rats, induces a better human obesity model than very high-fat diet | Journal Of Food Biochemistry | 2021 | 4 | Lima Tr;Voltarelli Fa;Freire Ls;Da Silva Fa;De Almeida Et;De Franca Sa;Pereira Mp;Damazo As;Navalta Ca;Kawashita Nh | WOS |
| 62 | Dha/epa supplementation decreases anxiety-like behaviour, but it does not ameliorate metabolic profile in obese male rats | British Journal Of Nutrition | 2022 | 4 | Neto J;Jantsch J;De Oliveira S;Braga Mf;Castro Lfds;Diniz Bf;Moreira Jcf;Giovenardi M;Porawski M;Guedes Rp | SCOPUS |

| | | | | | | |
|---|---|---|---|---|---|---|
| **63** | Enduring effects of an unhealthy diet during adolescence on systemic but not neurobehavioural measures in adult rats | Nutritional Neuroscience | 2022 | 4 | Nicolas S;Leime Cs;Hoban Ae;Hueston Cm;Cryan Jf;Nolan Ym | SCOPUS |
| **64** | A long-term energy-rich diet increases prefrontal bdnf in sprague-dawley rats | Nutrients | 2022 | 4 | Virtuoso A;Tveden-Nyborg P;Schou-Pedersen Amv;Lykkesfeldt J;Mueller B;Sorensen Db | WOS |
| **65** | Early postnatal exposure to a cafeteria diet interferes with recency and spatial memory, but not open field habituation in adolescent rats | Developmental Psychobiology | 2021 | 4 | Wait J;Burns C;Jones T;Harper E;Langley-Evans Sc;Voigt Jp | WOS |
| **66** | Caloric restriction or cafeteria diet from birth to adulthood increases the sensitivity to ephedrine in anxiety and locomotion in wistar rats | Physiology & Behavior | 2021 | 4 | Rocha-Gomes A;Teixeira Ae;Lima Dss;Rocha Lds;Da Silva Aa;Lessa Mr;Pinto Nad;Stuckert-Seixas Sr;Riul Tr | PUBMED |
| **67** | Restricted cafeteria feeding and treadmill exercise improved body composition, metabolic profile and exploratory behavior in obese male rats | Scientific Reports | 2022 | 3 | Alvarez-Monell A;Subias-Gusils A;Marine-Casado R;Belda X;Gagliano H;Pozo Oj;Boque N;Caimari A;Armario A;Solanas M;Escorihuela Rm | WOS |
| **68** | Short-term cafeteria diet is associated with fat mass accumulation, systemic and amygdala inflammation, and anxiety-like behavior in adult male wistar rats | Neuroscience | 2023 | 3 | Giovana Maciel Reis C;Rocha-Gomes A;Escobar Teixeira A;Gomes De Oliveira D;Mainy Oliveira Santiago C;Alves Da Silva A;Regina Riul T;De Jesus Oliveira E | SCOPUS |
| **69** | Palatable high-fat diet intake influences mnemonic and emotional aspects in female rats in an estrous cycle-dependent manner | Metabolic Brain Disease | 2021 | 3 | Silva Sp;Beserra-Filho Jia;Kubota Mc;Cardoso Gn;Freitas Frs;Goncalves Bsm;Vicente-Silva W;Silva-Martins S;Custodio-Silva Ac;Soares-Silva B;Maria-Macedo A;Santos Jr;Estadella D;Ribeiro Am | PUBMED |
| **70** | Behavioral and metabolic effects of a calorie-restricted cafeteria diet and oleuropein supplementation in obese male rats | Nutrients | 2021 | 3 | Subias-Gusils A;Alvarez-Monell A;Boque N;Caimari A;Del Bas Jm;Marine-Casado R;Solanas M;Escorihuela Rm | PUBMED |
| **71** | Anxiety associated with palatable food withdrawal is reversed by the selective faah inhibitor pf-3845: a regional analysis of the contribution of endocannabinoid signaling machinery | International Journal Of Eating Disorders | 2023 | 2 | De Ceglia M;Di Bonaventura A;Friuli M;Di Bonaventura Em;Gavito Al;Botticelli L;Gaetani S;De;Fonseca Fr;Cifani C | WOS |
| **72** | Early postoperative exposure to high-fat diet does not increase long-term weight loss or fat avoidance after roux-en-y gastric bypass in rats | Frontiers In Nutrition | 2022 | 2 | Ismaeil A;Gero D;Boyle Cn;Alceste D;Taha O;Spector Ac;Lutz M | WOS |
| **73** | Effects of omega-3 supplementation on anxiety-like behaviors and neuroinflammation in wistar rats following cafeteria diet-induced obesity | Nutritional Neuroscience | 2023 | 2 | Gonzalez Lpf;Da Silva Rodrigues J;De Farias Fraga G;Squizani Lf;Correia Ll;Neto Jp;Giovenardi M;Porawski M;Guedes Rp | WOS |
| **74** | Impact of cafeteria diet and n3 supplementation on the intestinal microbiota, fatty acids levels, neuroinflammatory markers and social memory in male rats | Physiology & Behavior | 2023 | 2 | Neto J;Jantsch J;Rodrigues F;Squizani S;Eller S;Oliveira Tf;Silveira Ak;Moreira Jcf;Giovenardi M;Porawski M;Guedes Rp | WOS |
| **75** | High-sugar/high-fat diet modulates the effects of chronic stress in cariocas high- and low-conditioned freezing rats | Physiology & Behavior | 2022 | 2 | Lages Yv;Maisonnette Ss;Rosseti Fp;Krahe Te;Landeira-Fernandez J | PUBMED |
| **76** | A laboratory environment previously associated with a palatable diet can result in overfeeding in rats | Indian Journal Of Animal Research | 2018 | 1 | Gabriela Martinez A;Lopez-Espinoza A;Josefina;Lopez-Uriarte P;Patricia Beltran-Miranda C;Daniel;Miguel-Gomez H;Cristina Espinoza-Gallardo A | WOS |

| | | | | | | |
|---|---|---|---|---|---|---|
| **77** | Antiobesity effect of polyherbal formulations in cafeteria and atherogenic diet induced obesity in rats | International Journal Of Pharmaceutical And Clinical Research | 2013 | 1 | Vikas Kumar J;Vishal B;Rajesh Kumar N | SCOPUS |
| **78** | Characterization of mitogen-activated protein kinase expression in nucleus accumbens and hippocampus of rats subjected to food selection in the cafeteria diet protocol | Cns & Neurological Disorders Drug Targets | 2016 | 1 | Sarro-Ramirez A;Sanchez D;Tejeda-Padron A;Buenfil-Canto Lv;Valladares-GarciaA J;Pacheco-Pantoja E;Arias-Carrion O;Murillo-Rodriguez E | PUBMED |
| **79** | Binge eating behavior and incentive motivation with a cafeteria diet | Behavioural Processes | 2021 | 1 | Vazquez-Herrera Nv;Zepeda-Ruiz Wa;Velazquez-Martinez Dn | PUBMED |
| **80** | Fat intake and obesity-related parameters predict striatal bdnf gene expression and dopamine metabolite levels in cafeteria diet-fed rats | Neuroscience | 2022 | 1 | Vindas-Smith R;Quesada D;Hernandez-Solano Mi;Castro M;Sequeira-Cordero A;Fornaguera J;Gomez G;Brenes Jc | PUBMED |
| **81** | Impact of calorie-restricted cafeteria diet and treadmill exercise on sweet taste in diet-induced obese female and male rats | Nutrients | 2023 | 0 | Alvarez-Monell A;Subias-Gusils A;Marine-Casado R;Boque N;Caimari A;Solanas M;Escorihuela Rm | SCOPUS |
| **82** | Extra virgin coconut oil (cocos nucifera l.) Intake shows neurobehavioural and intestinal health effects in obesity-induced rats | Food & Function | 2023 | 0 | Araujo De Vasconcelos Mh;Tavares Rl;Dutra Mldv;Batista Ks;D'oliveira Ab;Pinheiro Ro;Pereira Ra;Lima Mds;Salvadori Mgdss;De Souza El;Magnani M;Alves Af;Aquino Js | PUBMED |
| **83** | Transcranial direct current stimulation (tdcs) promotes state-dependent effects on neuroinflammatory and behavioral parameters in rats chronically exposed to stress and a hyper-palatable diet | Neurochemical Research | 2023 | 0 | De Castro Jm;De Freitas Js;Stein Dj;De Macedo Ic;Caumo W;Torres Ils | SCOPUS |
| **84** | Consumption of cashew nut induced anxiolytic-like behavior in dyslipidemic rats consuming a high fat diet | Behavioural Brain Research | 2023 | 0 | Dias Ccq;Madruga Ms;Almeida Gho;De Melo Mfft;Viera Vb;Bertozzo Ccms;Dutra Lmg;Alves Fa;Bezerra Jkb | WOS |
| **85** | Food selection of cafeteria diet affects memory dysfunction related to obesity | Neurochemical Research | 2019 | 0 | Feijo Gds;De Oliveira S;Thoen R;Schaab Ee;De Moura Ac;Franco F;Giovenardi M;Porawski M;Guedes Rp | PUBMED |

***Table S1. Total of articles included in the bibliometric analysis.*** *Were 85 the articles resulting from the applicability of criteria eligibility and filtering by duplicates. The table has the title of the article, journal, year of release, citations from 2013 to 2023, authors and database were it was found.*

| | Term | Strength | Rank |
|---|---|---|---|
| 1 | Obesity | 535 | 57 |
| 2 | Weight | 465 | 56 |
| 3 | Behavior | 441 | 55 |
| 4 | Metabolic | 387 | 54 |
| 5 | Palatable | 335 | 53 |
| 6 | Behavioral | 286 | 52 |
| 7 | Anxiety | 281 | 51 |
| 8 | Alterations | 220 | 50 |
| 9 | High-fat | 204 | 49 |
| 10 | Induced obesity | 194 | 48 |
| 11 | Memory | 193 | 47 |
| 12 | Stress | 191 | 45 |
| 13 | Treatment | 191 | 45 |
| 14 | Adiposity | 190 | 43 |
| 15 | Receptor | 190 | 43 |
| 16 | Hippocampus | 189 | 42 |
| 17 | Anxiety-like | 188 | 41 |
| 18 | Insulin | 180 | 40 |
| 19 | Health | 179 | 38 |
| 20 | Sugar | 179 | 38 |
| 21 | Glucose | 177 | 37 |
| 22 | Obese | 171 | 36 |
| 23 | Diet-induced obesity | 157 | 35 |
| 24 | Sucrose | 156 | 34 |
| 25 | Reward | 152 | 32 |
| 26 | Cholesterol | 152 | 32 |
| 27 | Behaviors | 145 | 31 |
| 28 | Anxiety-like behavior | 138 | 30 |
| 29 | Disorders | 130 | 29 |
| 30 | Inflammation | 127 | 27 |
| 31 | Cortex | 127 | 27 |
| 32 | Hedonic | 126 | 26 |
| 33 | Caloric intake | 112 | 25 |
| 34 | Energy intake | 109 | 24 |
| 35 | Recognition | 108 | 23 |
| 36 | Highly palatable | 107 | 22 |
| 37 | Cognitive | 104 | 19 |
| 38 | Learning | 104 | 19 |
| 39 | Palatable cafeteria | 104 | 19 |
| 40 | Metabolic syndrome | 100 | 18 |
| 41 | Accumbens | 87 | 16 |
| 42 | Nucleus accumbens | 87 | 16 |
| 43 | Dopamine | 85 | 15 |
| 44 | Obesogenic | 82 | 14 |
| 45 | Cafeteria-style | 62 | 13 |
| 46 | Amygdala | 52 | 12 |
| 47 | Neuroinflammatory | 50 | 11 |
| 48 | Cafeteria-diet | 47 | 9 |
| 49 | Cafeteria diet-induced | 47 | 9 |
| 50 | Hypercaloric | 46 | 8 |
| 51 | Microbiota | 45 | 7 |
| 52 | Anxiety-like behaviour | 37 | 6 |
| 53 | Spatial memory | 35 | 5 |
| 54 | Metabolic profile | 32 | 4 |
| 55 | Intestinal | 27 | 3 |
| 56 | Neurobehavioural | 23 | 2 |
| 57 | Stress-induced | 22 | 1 |

***Table S2. Terms of the document feature matrix.*** *These terms were weighted by their occurrence and then fit to a matrix that rank them. The 1st position of the rank is the less weight term.*

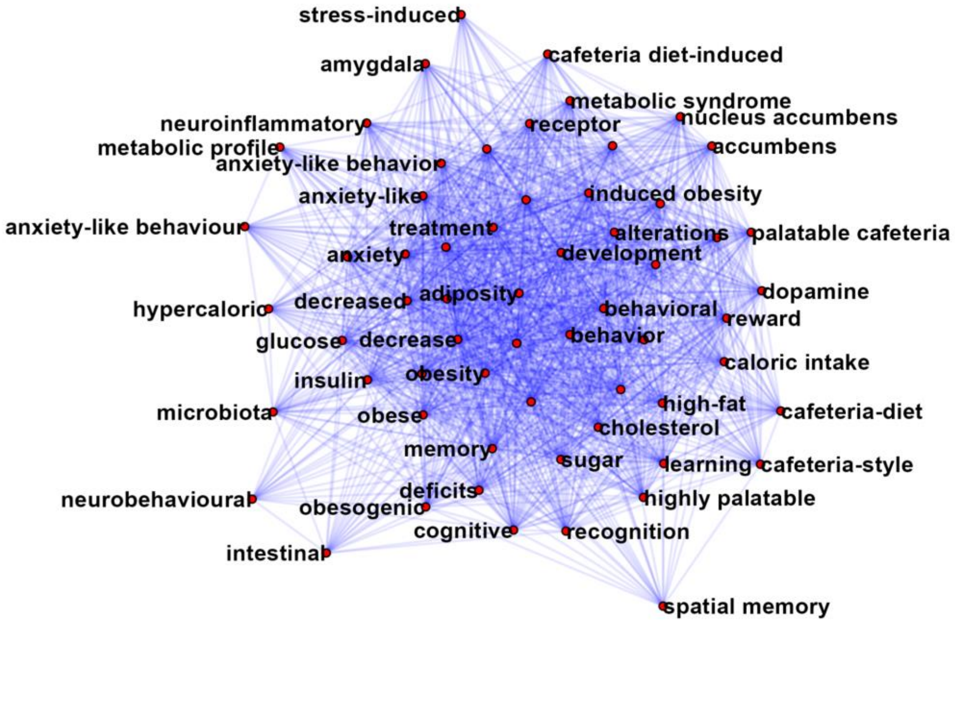


***Fig S1. Strength and weights network of the terms.*** *A preliminary network modeling of the terms shown the relationship of the terms within them.*
